# Trimethylamine *N*-oxide reduces neurite density and plaque intensity in a murine model of Alzheimer’s disease

**DOI:** 10.1101/2022.05.15.492014

**Authors:** Katie R. Zarbock, Jessica H. Han, Ajay P. Singh, Sydney P. Thomas, Barbara B. Bendlin, John M. Denu, John-Paul J. Yu, Federico E. Rey, Tyler K. Ulland

## Abstract

**Background:** Alzheimer’s disease (AD) is the most common aging-associated neurodegenerative disease; nevertheless, the etiology and progression of the disease is still incompletely understood. We have previously shown that the microbially-derived metabolite trimethylamine *N*-oxide (TMAO) is elevated in the cerebrospinal fluid (CSF) of individuals with cognitive impairment due to AD and positively correlates with increases in CSF biomarkers for tangle, plaque, and neuronal pathology.

**Objective:** We assessed the direct impact of TMAO on AD progression.

**Methods:** To do so, transgenic 5XFAD mice were supplemented with TMAO for 12 weeks.

**Results:** Oral TMAO administration resulted in significantly reduced neurite density in several regions of the brain, as assessed through quantitative brain microstructure imaging with neurite orientation dispersion and density imaging (NODDI) magnetic resonance imaging (MRI). Amyloid-β plaque mean intensity was reduced, while plaque count and size remained unaltered. Proteomics analysis of the cortex revealed that TMAO treatment impacted the expression of 31 proteins (1.5-fold cut-off) in 5XFAD mice, including proteins known to influence neuronal health and amyloid-β precursor protein processing. TMAO treatment did not alter astrocyte and microglial response (as determined by histological analysis) nor cortical synaptic protein expression.

**Conclusion:** These data suggest that elevated plasma TMAO impacts AD pathology via reductions in neurite density.

## INTRODUCTION

Alzheimer’s disease (AD), leading to cognitive decline and dementia, is characterized by two key histopathological hallmarks: the presence of extracellular plaques composed primarily of amyloid-β (Aβ) peptides generated from the cleavage of amyloid-β precursor protein (AβPP) and intraneuronal neurofibrillary tau tangles composed of hyperphosphorylated tau. These protein aggregates lead to synaptic and neuronal damage and trigger microglial and astrocyte activation. Although AD has been extensively studied, its etiology and progression are still not fully understood.

In recent years the gut microbiome has emerged as a potential modifier of AD progression. Animal studies showed that APPPS1 germ-free mice had reduced Aβ pathology in comparison to specific pathogen free conventionally-raised mice, suggesting that the gut microbiota influence disease progression [1]. Treatment of APPswe/PS1ΔE9 mice with antibiotics throughout life [2] and for one week beginning at postnatal development day 14 [3] significantly reduced plaque burden and neuroinflammation. Examining the gut microbiome in humans with AD, our team showed that individuals with AD have significant alterations in the abundance of several bacterial taxa relative to healthy controls [4]. Furthermore, we found that cerebrospinal fluid (CSF) levels of bacterial-derived trimethylamine *N*-oxide (TMAO) were elevated in human individuals with mild cognitive impairment and dementia due to AD as compared to their cognitively unimpaired controls [5]; levels of CSF TMAO also positively correlated with CSF biomarkers for tangle pathology alone (phosphorylated tau [p-tau]), tangle and plaque pathology (p-tau:Aβ42), and neuronal damage (total tau and neurofilament light chain).

TMAO is generated primarily from choline and L-carnitine through bacterial and host metabolic pathways; first, bacteria convert these molecules into TMA [6–7], which is then absorbed and transported to the liver where flavin-containing monooxygenases (FMOs), primarily FMO3, convert it into the soluble, polar molecule TMAO [8]. This metabolite then enters vascular circulation – where it is detected at micromolar concentrations – eventually reaches the brain, and is cleared from the body primarily via the kidneys [9–10].

TMAO has been shown to worsen diseases occurring in highly vascularized organs, including non-alcoholic fatty liver disease, thrombosis, atherosclerosis, renal disease, and hippocampal senescence [11–15]. TMAO levels are higher in plasma [16] and brain tissue [16–17] of aging mice and are also elevated in plasma and CSF of elderly humans and humans with AD, respectively [5,16]. TMAO levels in plasma and brain tissue are elevated in 3xTg-AD mice, a model of AD [17]. High choline- and TMAO-supplemented WT mice displayed elevated plasma TMAO levels and decreased cognitive function [16, 18]. However, to the best of our knowledge, no study to date has investigated the effect of elevated levels of plasma TMAO on AD pathology *in vivo*. To address this question, we administered TMAO to transgenic 5XFAD female mice and examined synaptic, neurite, and amyloid-β plaque pathology, the cortical proteome, and the glial response. Our results suggest that TMAO reduces neurite density and decreases amyloid-β plaque intensity.

## MATERIALS AND METHODS

| Antibodies and Stains | Company | Product ID |
| --- | --- | --- |
| β-actin, Rabbit | Cell Signaling Technologies | 4967S |
| GFAP, Rabbit pAb | Invitrogen | 14-9892-82 |
| Anti-Goat, Donkey pAb | Abcam | ab150129 |
| Anti Iba-1, Goat pAb | Abcam | ab5076 |
| IRDye 680RD Goat anti-mouse | LI-COR | 926-68070 |
| IRDye 800CW Donkey anti-Rabbit | LI-COR | 926-32213 |
| Methoxy-X04 | Tocris Bioscience | 4920 |
| PSD95, Rabbit mAb | Cell Signaling Technologies | 3409S |
| Synaptophysin, Mouse mAb | Agilent | M731529-2 |
| TOPRO-3 Iodide | Invitrogen | T3605 |

### Mice

All mice were handled as outlined in animal use protocols approved by the University of Wisconsin-Madison Animal Care and Use Committee. 5XFAD mice were purchased from The Jackson Laboratory and bred in the University of Wisconsin-Madison Biomedical Research Models Services specific pathogen free facility; female pups were evenly split from each litter between the two groups, using hemizygote females (untreated group, n=11; TMAO-treated group, n=10) and WT female littermate controls (untreated group, n=5; TMAO-treated group, n=5). Mice were housed on a 12-hour light/dark cycle and given *ad libitum* access to rodent chow. Regular water was provided to breeders and pups. At 5-6 weeks of age, mice either remained on regular drinking water or were placed on 0.3%w/v TMAO drinking water. TMAO was administered *ad libitum* for approximately 12 weeks; mice were sacrificed at 4 months of age.

Mice were anesthetized with isoflurane and underwent cardiac puncture for a terminal blood draw, followed by cervical dislocation. The brain was removed, placed into ice-cold PBS, and sagittally bisected. The right hemisphere was fixed in 4% paraformaldehyde (PFA) and prepared for histology, as described below; the left hemisphere was dissected out into the cortex, hippocampus, and cerebellum, and tissue was snap frozen in liquid nitrogen. Blood was spun down at 3,000 x *g* for 15 minutes at 4°C; the resulting plasma layer was removed and snap frozen. All snap frozen tissues were stored at −80°C until needed. Sacrifices were performed between 1:00 and 4:00 PM.

### TMAO plasma quantification

Plasma TMAO levels were measured by LC-MS/MS. One volume of plasma and 4 volumes of ice-cold extraction solution (HPLC-grade MeOH with 2.5 μM d9-TMAO internal control) were combined and spun at 21,100 x *g* (maximum speed) for 3 minutes at 4°C. One volume of the resulting supernatant was then mixed with 1 volume HPLC-grade water. The resulting samples were injected onto a Waters ACQUITY C18 UPLC column (1.7 μm, 2.1 mm x 100 mm) that was coupled to a Thermo Fisher Q-Exactive mass spectrometer at a flow rate of 0.2 mL/min. Elution of samples occurred over a 7-minute isocratic gradient of 25% water + 5 mM ammonium acetate + 0.05% acetic acid and 75% methanol. TMAO was quantified in the positive mode using Parallel Reaction Monitoring (PRM) using an inclusion list of 76.076 and 85.132 m/z (for TMAO and d9TMAO respectively). El-MAVEN was used for peak quantification, and internal standards were utilized as a comparison to calculate plasma concentrations.

### Western blot analysis

Total protein was extracted from the snap frozen cortex of the left hemisphere via Dounce homogenization in RIPA buffer from control (n=6) or TMAO-treated (n=4) mice. Protein extracts were quantified using BCA and diluted to 2 μg/μL with water and the appropriate amount of 4x Laemmli buffer and 2-mercaptoethanol. 15 μg of protein per sample was loaded into individual wells of a 4% - 20% Mini-PROTEAN TGX precast protein gel and ran at 200V for 25 minutes using a Bio-Rad Mini-PROTEAN gel system; tris-glycine SDS running buffer was used. Protein was transferred to a methanol-soaked PVDF membrane at 100V for 1 hour in tris-glycine transfer buffer with 20% methanol using a Bio-Rad Mini Trans-Blot module. The membrane was blocked in 5% nonfat-dry milk (blotting-grade blocker) in TBS. Following blocking, PSD-95 (1:1000), synaptophysin (1:1600), β-actin (1:500), and Tween-20 (0.1%) were added directly to the blocking buffer, and the membrane was incubated on a rocking platform a 4°C for two nights. After ~40 hours, the membrane was washed 5 times with TBST and then incubated in the dark with 800CW anti-rabbit and 680RD anti-mouse antibodies (1:15,000), rocking at room temperature for 1 hour. After incubation with the secondary antibodies, the membrane was washed 5 times with TBST and 2 times with TBS. The membrane was imaged using a LI-COR Odyssey instrument and was exposed for 2 minutes for both the 700 and 800 channels. The resulting image was analyzed in LI-COR Image Studio Lite (version 5.2), and the signal of the synaptic band was normalized to the signal of the β-actin band for each sample.

### Ex vivo *MRI*

Mice receiving control (n=5) or TMAO (n=4) drinking water between 4 and 5 months of age (mean ± STD: 4.516 ± 0.411 months) were anesthetized and transcardially perfused with ice-cold PBS and 4% PFA; brains were then cleanly dissected *en bloc* from the cranial vault. *Ex vivo* imaging and analysis, including standard data preprocessing, and study template generation, was performed as previously described [19]. Briefly, multislice, diffusion-weighted spin echo images were used to acquire 10 non-diffusion-weighted images (b=0 s/mm^2^) and 75 diffusion-weighted images (25 images: b=800 s/mm^2^; 50 images: b=2000 s/mm^2^) using noncollinear, diffusion-weighting directions. Diffusion imaging was performed with an echo time of 24.17/2000 ms, field of view = 30×30 mm^2^, and matrix = 192×192 reconstructed to 256×256 for an isotropic voxel size of 0.25 mm over 2 signal averages. Multi-shell diffusion data were fit with the Microstructure Diffusion Toolbox [20] to the NODDI *ex vivo* model. An additional compartment of isotropic restriction was included to account for potential fixative effects as recommended [21]. Regions of interest (ROIs) were selected *a priori* and defined with a DTI-based mouse brain atlas [22]. A Student’s *t-*test was used to determine the significance (*p* < 0.05) of ROI values between female 5XFAD mice receiving regular water and TMAO supplementation; statistically significant differences were determined after controlling for multiple comparisons with the Benjamini–Hochberg procedure (false discovery rate = 0.05).

### Shotgun proteomics

Approximately 50 mg of tissue from the cortex was Dounce homogenized and tip-sonicated in 0.5% SDS/Tris-HCl. Protein was quantified by BCA and 50 μg was used for the shotgun proteomics. Samples were denatured and alkylated by addition of 10 mM dithiothreitol (DTT) and 20 mM iodoacetamide (IAA), respectively. Any remaining detergent was removed using a chloroform methanol extraction. The concentrated protein samples were reconstituted and tip-sonicated in 2M urea/NH4HCO3. Then the samples were digested with trypsin overnight, de-salted and cleaned with C18 stage tips, dried down, and resuspended in sample diluent (5% ACN, 0.1% acetic acid in HPLC-grade water).

The separated peptides were then injected onto the same LC-MS/MS system as used for plasma TMAO quantification. Mobile phase consisted of water + 0.1% formic acid (A) and acetonitrile + 0.1% formic acid (B). Resolution of peptides was performed with a 2-step linear gradient of 2% to 35% B over 90 minutes followed by 35% to 95% B over 12 minutes. Data-dependent acquisition (DDA) mode was utilized for all samples.

MaxQuant (Max Planck Institute) was applied to process library samples. Identification and quantification of proteins were performed using Perseus with the MaxQuant library, a *Mus musculus* reference genome FASTA (SWISS-PROT canonical), and default settings. Differential expression of proteins was considered significant if q < 0.05.

### Brain sample preparation for histology

The right hemisphere of the brain was fixed in 4% PFA for 48 hours, rinsed with PBS, and then cryoprotected in 30% sucrose for 48 hours. Brains were then frozen in a mixture of 2 parts 30% sucrose and 1 part OCT compound. Serial 40-μm coronal brain sections were cut on a freezing sliding microtome from the rostral anterior commissure to the caudal hippocampus.

### Histology preparation, imaging, and quantification

Three sections spaced 440 μm apart from one another (Bregma −1.42 to −2.3 mm) were stained for plaques (methoxy-X04), nuclei (TOPRO-3), and either astrocytes (GFAP) or microglia (Iba-1).

To assess plaque burden, a micrograph of the entire section was captured in a low-powered field at a single focal plane using a Nikon A1R+ confocal microscope; the same power, gain, and pinhole settings were used for each channel for every image captured. The percent area positive for methoxy-X04 in the cortex and the hippocampus was quantified using FIJI. To assess plaque count, size, and mean intensity, high-powered field z-stack images were captured (447 x 447 x 40 μm; 0.7 μm thickness) in the retrosplenial and somatosensory areas of the cortex. Images were processed using Imaris software, with the Surface function applied to assess plaques. The surfaces were split using an automated Matlab script, and the number of resulting objects was used to determine plaque count. The mean volume of the split plaque surfaces was used to calculate group mean plaque volume. The average value for mean intensity for the plaque surfaces was used to quantify plaque intensity.

To assess the total number of immune cells and immune cells surrounding a plaque, the Imaris Spot function was utilized. The astrocyte or microglia channel was colocalized with the nuclear channel, and a separate colocalization channel was generated. Spots 8 μm in diameter and 6 μm in diameter were generated using this colocalization channel to label astrocytes and microglia, respectively. Spots were quality controlled by reviewing the morphology of the immune cell that the Spot function identified, and any errant spots were removed. The number of spots was used to determine the total number of immune cells. An automated surface-spot distance calculating Matlab script was used to generate the number of immune cells within 15 and 30 μm of a plaque.

For each metric, the resulting values for the 3 sections were averaged for each animal, and animal averages were used to generate a group mean for the retrosplenial and the somatosensory regions. To calculate the “averaged” data, all 6 images per animal (i.e., the 3 retrosplenial images and the 3 somatosensory images) were averaged, and the animal averages were used to calculate a group mean.

## RESULTS

### Elevated plasma TMAO leads to reduced neurite density throughout the brain

To assess the impact of TMAO on the development of AD pathology, we administered 0.3% w/v TMAO in drinking water, *ad libitum*, to 5XFAD female mice for 12 weeks starting at 5-6 weeks of age; control animals consumed regular water (Fig. 1A). Mice were sacrificed at 4 months of age, as assessing phenotypes at this time point allowed for analysis of the initial impacts of TMAO on AD pathology. Liquid chromatography tandem mass spectrophotometry (LC-MS/MS) analysis of plasma samples collected at the end of treatment showed that TMAO was significantly increased in the TMAO-administered 5XFAD mice relative to their control counterparts (Fig. 1B); similar differences were observed in WT mice (supplementary fig. 1A).

**Figure 1.**
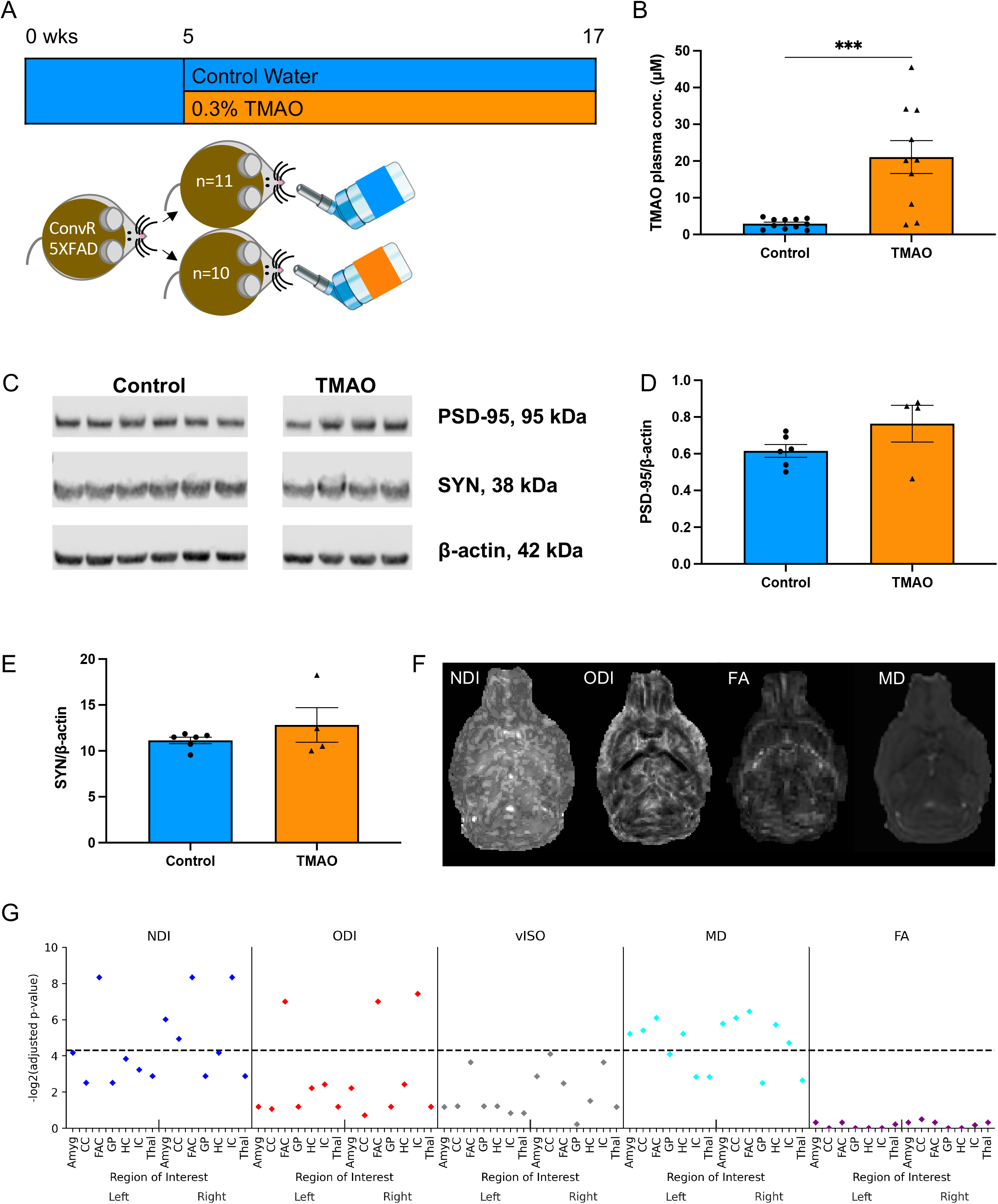
Effects of TMAO supplementation on synaptic protein expression and neurite density. (A) 5XFAD female mice received either control, regular water (control, n=11) or water supplemented with 0.3% TMAO (TMAO, n=10). (B) Plasma TMAO levels were significantly increased in animals receiving TMAO. (C) Representative blots showing expression of postsynaptic density-95 (PSD-95), synaptophysin (SYN), and β-actin. Total protein was isolated from the cortex of control (n=6) and TMAO-treated (n=4) mice. Normalized protein signals of PSD-95 (D) and SYN (E) were quantified. Data in B, D, and E are shown as the mean ± SEM (two sample t-test); *** denotes p < 0.001. (F) Mice receiving control (n=5) or TMAO (n=4) drinking water were sacrificed between 4 and 5 months, and *ex vivo* MRI brain imaging was performed. NODDI, or Neurite Orientation Dispersion and Density Imaging, (NDI [neurite density index], ODI [orientation dispersion index], vISO [isotropic volume]) and DTI (FA [fractional anisotropy], MD [mean diffusivity]) were completed (axial images shown). (G) Several regions of interest from both hemispheres were assessed: amygdala (Amyg), corpus callosum (CC), frontal association cortex (FAC), globus pallidus (GP), hippocampus (HC), internal capsule (IC), and thalamus (Thal). P-values (-log2 transformed) are displayed on the y-axis; the horizontal dashed line denotes the FDR < 0.05 cutoff with values below the line showing no statistical significance.

Because our clinical analysis revealed significant positive correlations between CSF TMAO levels and biomarkers of neuronal damage [5], we determined the impact of TMAO on neuronal health. First, synaptic protein expression was examined. We extracted total protein from the entire left cortex and performed western blot analysis to assess the expression of presynaptic protein synaptophysin and postsynaptic protein postsynaptic density-95 (PSD-95). There was no change in the normalized expression of these proteins between control and TMAO-treated 5XFAD mice (Fig. 1C-E).

We next used the more sensitive technique of diffusion weighted imaging to assess neurite density in 5XFAD mice (treated as described above; control, n=5; TMAO-treated, n=4). *Ex vivo* multi-compartment diffusion weighted imaging was performed. Imaging data were fit to neurite orientation dispersion and density imaging (NODDI) (encompassing neurite density [NDI], neurite orientation dispersion [ODI], and isotropic volume [vISO]) and diffusion tensor imaging (DTI) (encompassing mean diffusivity [MD] and fractional anisotropy [FA]) models; region-specific changes were assessed. TMAO-treated animals had significantly reduced NDI in bilateral frontal association cortex, right amygdala, corpus callosum, and the internal capsule (Fig. 1F-G). ODI was also significantly reduced in TMAO-treated mice in the bilateral frontal association cortex and internal capsule. In addition, MD was also significantly reduced in bilateral amygdala, corpus callosum, frontal association cortex, hippocampus, and in the right internal capsule in TMAO-treated mice. FA changes were not observed in any region of the brain. These results indicate that elevated levels of plasma TMAO lead to concomitant reductions in neurite density and dispersion.

### TMAO reduces plaque mean intensity without major changes to plaque burden

Assessment of plaque pathology (Fig. 2A) disclosed that TMAO did not alter the plaque burden in the hippocampus, although there was a trend towards reduced plaque area in the cortex (Fig. 2B; p = 0.088); the same trend towards reduced plaque area in the cortex was observed in gnotobiotic mice (n=4/group) colonized with defined bacterial synthetic communities that differed in their capacity to metabolize choline and accumulate TMA/TMAO (Fig. S1; p = 0.059).

**Figure 2.**
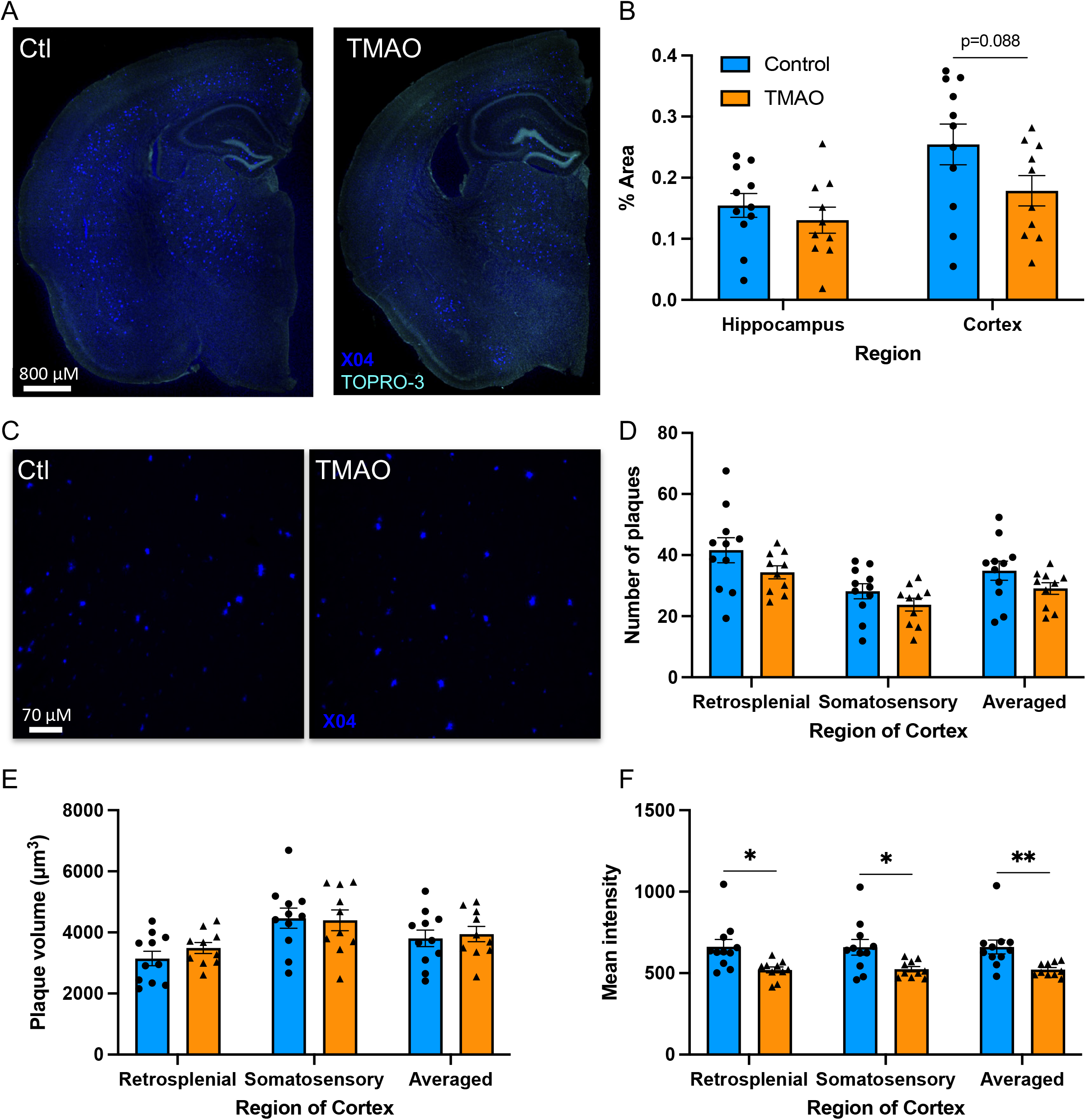
TMAO reduces plaque intensity but does not influence amyloid plaque burden. Brain sections were stained for plaques (methoxy-X04, dark blue) (A, C) and nuclei (TOPRO-3, turquoise blue) (A). Percent area positive for methoxy-X04 in the cortex and the hippocampus (B). High-powered field, confocal images of the retrosplenial and somatosensory regions of the cortex (C). Number of plaques/field (D), volume/plaque (E), and mean intensity/plaque (F). Three sections were quantified and averaged for each animal. Data are shown as the mean ± SEM (n=11 control, n=10 TMAO, two sample t-test). * denotes p < 0.05 and ** denotes p < 0.01.

TMAO did not impact the number or size of plaques in the cortex (Fig. 2C-E) but significantly reduced the intensity of plaques in both the retrosplenial and somatosensory regions of the cortex (Fig. 2F). Taken together, these data indicate that TMAO administration had a minimal impact on amyloid-β plaque pathology in 5XFAD mice.

### TMAO alters the cortex proteome

To clarify the complex responses of the brain to TMAO (i.e., significant reduction of neurite density with minimal influence on plaques), we performed a quantitative proteomic analysis of the cortex in WT and 5XFAD mice supplemented with TMAO and control animals (n=5-6/group). The entire cortex from one hemisphere of each mouse was homogenized, protein was extracted, and peptide libraries were prepared for shotgun proteomics using label-free quantitation methods. The resulting proteomics data was aligned to a reference database (*Mus musculus* genome FASTA [SWISS-PROT canonical]) and abundances (TMAO-treated *vs*. untreated) were calculated. Only a single protein (MIF [macrophage migration inhibitory factor]; −4.9-fold differentially expressed [DE]) was significantly regulated (q < 0.05) in TMAO-supplemented WT mice compared to untreated counterparts (supplemental fig. 1B), whereas 127 proteins were significantly DE when comparing TMAO-treated 5XFAD to their untreated 5XFAD counterparts (56 upregulated; 71 downregulated), 31 of which changed >1.5-fold (16 upregulated; 15 downregulated) (Fig. 3). Of particular interest, MIF (−5.7-fold DE), SYNJ1 (synaptojanin-1; 1.5-fold DE), and SYT1 (synaptotagmin-1; 1.6-fold DE) were among these 31 proteins. MIF, a pro-inflammatory cytokine, and SYNJ1, a phosphoinositide phosphatase localized in synapses and neuronal projections, both influence cognitive function *in vivo* and neuronal cell responses to Aβ oligomers *in vitro* [23–24]. SYT1, a calcium sensor involved in neurotransmitter vesicle release, interacts with and impacts proteins involved in AβPP processing, specifically presenilin-1 (PS1) and BACE1 (β-site AβPP cleaving enzyme 1), influencing Aβ peptide generation [25]. Altogether, these results, combined with the observed reduction in neurite density, suggest that TMAO administration alters the 5XFAD murine cortex proteome in a manner that promotes neurodegeneration.

**Figure 3.**
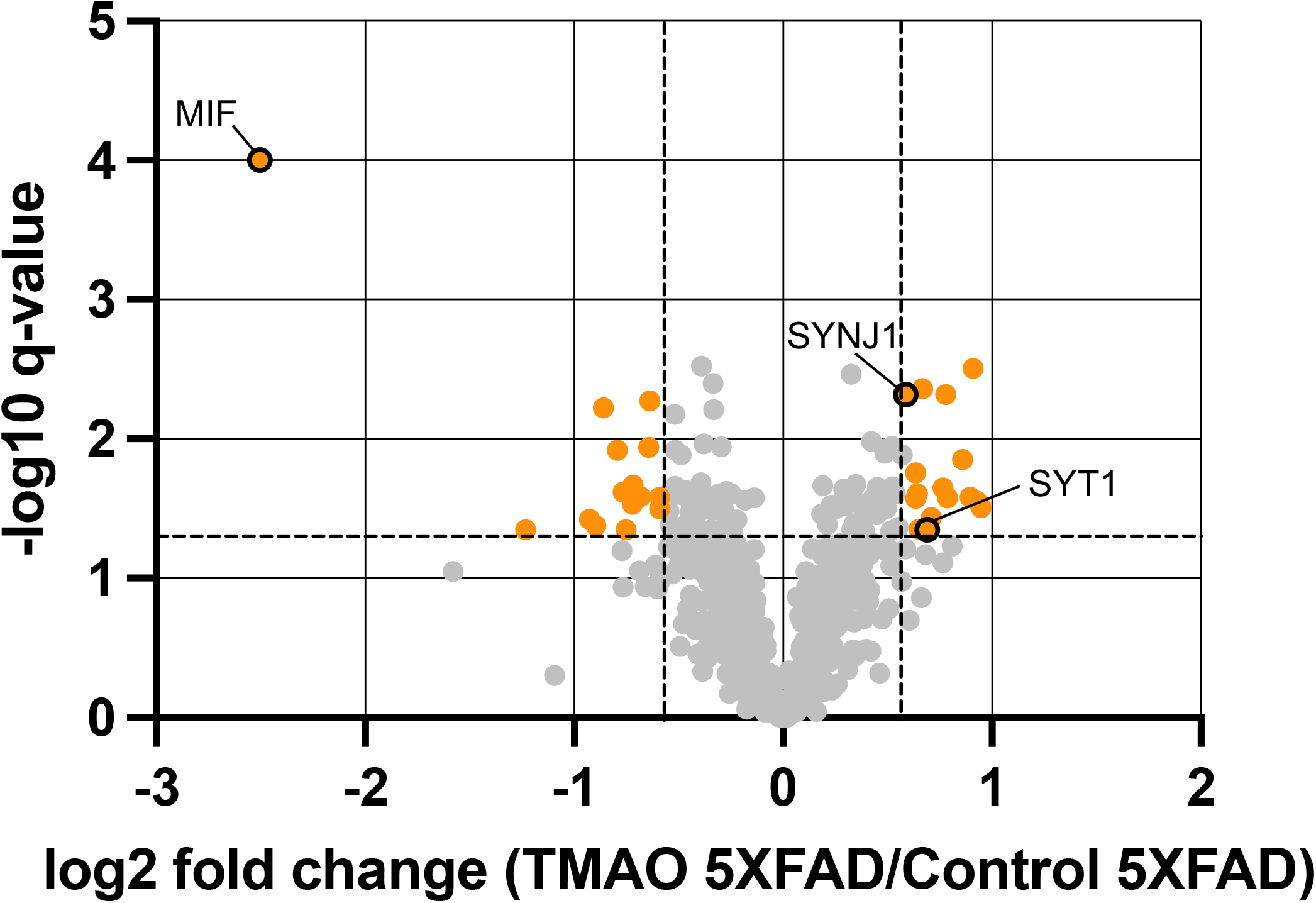
TMAO alters the cortex proteome of 5XFAD mice. Volcano plot showing log2 fold change (FC) of protein abundances of TMAO mice compared to those of control mice (n=5-6/group). The horizontal dashed line represents q = 0.05 and the vertical dashed lines represent FC = |1.5|. Proteins with q < 0.05 and FC > |1.5| are shown in orange, and proteins of interest are circled with a black border.

### TMAO does not impact the glial response

As MIF is a pro-inflammatory cytokine and was the most altered protein of the 5XFAD cortex proteome, we further assessed the glial response using histology. Using confocal microscopy, we quantified the total number of GFAP+ astrocytes per high-powered field and the number of GFAP+ cells within 15 and 30 μm of a plaque (Fig. 4A). We observed that, in TMAO-treated mice, the total number of astrocytes in the retrosplenial area of the cortex trended lower (Fig. 4B; p = 0.1); however, there was no observed difference in astrocyte number in the somatosensory region of the cortex or when both regions were averaged (Fig. 4B). The number of astrocytes surrounding a plaque was unaltered between groups (Fig. 4C).

**Figure 4.**
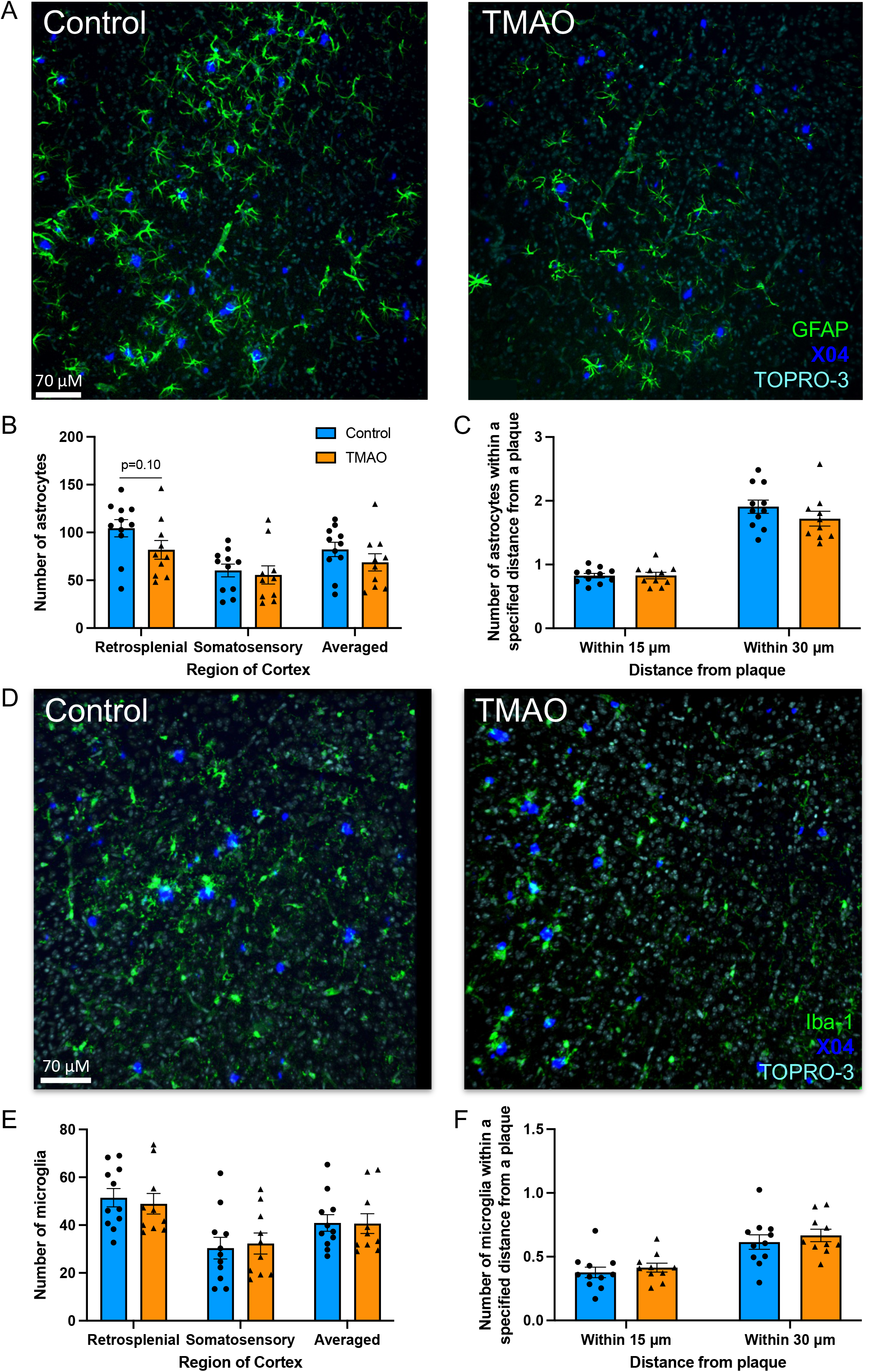
TMAO does not influence the glial response. Brain sections were stained for plaques (methoxy-X04, dark blue), nuclei (TOPRO-3, turquoise blue), and astrocytes (A) (glial acidic fibrillary protein [GFAP], green) or microglia (D) (ionized calcium-binding adaptor molecule 1 [Iba-1], green). Total number of astrocytes/field (B) or microglia/field (E). Number of astrocytes (C) or microglia (F) within 15 and 30 μm of a plaque (average of retrosplenial and somatosensory regions). Three sections were quantified and averaged for each animal. Data are shown as the group mean ± SEM (n=11 control, n=10 TMAO, two sample t-test).

Additionally, we determined the number of Iba-1+ microglia per high-powered field and within 15 and 30 μm of a plaque. TMAO did not significantly alter either the total number of microglia per high-powered field or the number of microglia surrounding plaques (Fig. 4D-F).

## DISCUSSION

TMAO has been shown to be elevated in both mice and humans during normal aging and AD [5, 16, 17], and positively correlates with CSF AD biomarkers [5]. We examined whether TMAO impacts AD pathology by supplementing 5XFAD female mice with TMAO in their drinking water. Our results suggest that TMAO reduces neurite density, neurite dispersion, and amyloid-β plaque intensity without changing other aspects of amyloid-β pathology, synaptic protein expression, or neuroimmune response.

Plasma TMAO concentrations were significantly higher in mice administered TMAO than in their control counterparts (21.1 vs 2.9 μM, Fig. 1B). In work examining the impact of TMAO on cardiac events, humans with and without such events were shown to have plasma TMAO levels as high as 312 μM; the interquartile range of the participants was 2.4 - 6.2 μM [12, 26]. The levels of TMAO in the current study fall within observed physiological ranges but are approximately three-fold greater than the upper value of the interquartile range of this previous work [12, 26]. However, TMAO levels observed in our study were lower than those attained in previous studies assessing the impact of this metabolite on cognition [16,18]. These interventions resulted in elevated levels of nonfasting plasma TMAO (approximately 30 and 90 μM, respectively) and cognitive deficits, as assessed by nest building, novel object recognition, Morris water maze, and shuttle box tests. Because cognitive decline is associated with neuronal pathology, it would be interesting to test cognitive capacity in our TMAO-treated 5XFAD mice in future studies.

To assess neuronal health in our study, we examined both synaptic protein expression via western blot and neurite density via *ex vivo* NODDI MRI. A reduction in synaptic protein expression was not observed at the age of sacrifice (Fig. 1C-E), suggesting that elevated TMAO levels do not influence synaptic protein expression at this relatively early disease stage. A previous study [15] found that 16 weeks of administration of 1.5% w/v TMAO drinking water to 24-week-old senescent-prone and -resistant male mice reduced the hippocampal expression of synaptic proteins synaptophysin, PSD-95, and N-methyl-d-aspartate 1 [15]. The discrepancy between the synaptic protein expression results of the two studies could be due to differences in TMAO concentration (previous study used 5X higher concentration), age at time of treatment, and duration of the treatment. These differences may have allowed for heightened and/or extended impact of TMAO on synapses in the earlier study [15].

While synaptic protein expression was unaltered, neurite density and dispersion were significantly reduced in mice with higher TMAO levels (Fig. 1F-G). NDI, ODI, and MD were significantly reduced in several regions of the brain, bilaterally, indicative of extensive detrimental effects of TMAO on neural microstructure. This includes the frontal association cortex, a region strongly impacted by plaque deposition in the 5XFAD model. FA was unaltered between groups, which represents the limits of the analysis capacity of DTI as compared to NODDI. This is likely because DTI relies on assessment of only a single compartment (intra-neuronal space), whereas NODDI assesses the extracellular compartment in addition to the intracellular compartment and is thus a more sensitive technique to assess neuropathological changes. The discrepancy between the synaptic protein expression and neurite density results might be explained by differences in sensitivity between methods, as the assessment of global synaptic protein expression in cortex homogenate from a large region of the brain is likely not as sensitive as *ex vivo* MRI, which has the capability to assess individual neurites and specific brain regions of interest. This is especially relevant at the young age and relatively early disease stage at which mice were sacrificed.

TMAO treatment resulted in no change in plaque number or size but in a significant reduction in plaque intensity and in a trend towards reduced cortical plaque burden (Fig. 2B, D-F). The current literature assessing the impact of TMAO on amyloid pathology shows similarly mixed results. One *in vitro* study demonstrated that activity of one of the enzymes leading to Aβ peptide formation - γ-secretase - was lowered by TMAO [27]. However, TMAO can also promote protein stabilization of the Aβ peptide and the conformational transformation of fibrils into β-pleated sheets, a necessary step in plaque formation [28, 29]. These *in vitro* studies, however, were conducted using supraphysiological concentrations of TMAO. Overall, TMAO had a very limited impact on plaque pathology in our study.

Several proteins regulated by TMAO have documented connections with AD, including MIF, SYNJ1, and SYT1. MIF has been shown to be elevated in brain tissue and CSF of humans with AD [23, 30] and in brain tissues of the APP23 transgenic AD mouse model [23]. APP23/MIF^+/-^ mice showed reduced memory and spatial learning in the Morris water maze test as compared to APP23 mice with WT levels of MIF, suggesting that a reduction in MIF expression in APP23 mice promotes cognitive impairment. Overexpression of MIF in SH-SY5Y neuroblastoma cells supported higher cell viability after exposure to Aβ oligomers than that of SH-SY5Y cells with WT MIF expression. These data suggest that MIF plays a neuroprotective role, and the reduction of MIF in the TMAO-treated mice may contribute to the reduction in neurite density.

SYNJ1 mRNA levels have been shown to be increased in the isocortex of humans with AD, particularly in APOEε4 carriers [31], and SNPs in the gene encoding for SNYJ1 correlate with AD cognitive impairment age of onset and cognitive testing scores [32]. SYNJ1 overexpression in C57BL/6 mice leads to increased cognitive impairment in aged mice as compared to their WT counterparts, as evaluated by fear conditioning and radial arm water maze tests [32], whereas reduced SYNJ1 expression in Tg2576 AD mice rescues cognitive deficits, as assessed by fear conditioning, radial arm water maze, and novel object recognition tests [24]. Additionally, Aβ peptides do less damage to dendritic spines in SYNJ1 knockout neuronal cultures [24], suggesting that reduction in SYNJ1 is protective to spine density in the presence of Aβ. These findings support the idea that an increase in SYNJ1 may have a negative effect on molecular and cognitive aspects of AD and may contribute to the observed reduction in neurite density.

SYT1, which was increased by TMAO in 5XFAD mice, influences proteins involved in APP processing and has been detected at higher levels in individuals with mild cognitive impairment and dementia due to AD [33]. SYT1 has been shown to interact with presenilin-1 (PS1) in dendrites in a calcium-dependent manner and influences the intracellular localization of PS1 and BACE1, as well as the conformation of PS1. Both PS1 and BACE1 are involved in the generation of Aβ peptides from AβPP, and with the influence SYT1 has on these two proteins, it is unsurprising that SYT1 knockdown led to decreased Aβ peptide levels and its overexpression led to increased levels [25]. SYT1 influences may be detected with increased length of TMAO treatment and age of mice at the time of sacrifice.

Overall, the proteomics data indicate that TMAO induces a number of important molecular changes in the cortices of mice, including several proteins previously implicated in AD progression. Importantly, these results also suggest that the effects of TMAO on the cortex proteome are modulated by AD-related phenotypes presented in 5XFAD mice, as there were more drastic alterations in response to TMAO in these animals than in WT mice subjected to the same treatment.

In summary, our study suggests that TMAO influences neuronal response to plaques resulting in neurite density and dispersion reduction. These reductions are relevant, as damage to neurons is highly correlated with cognitive impairment. While synaptic degradation was not detected via western blot, the MRI results may indicate that neurite density reduction was at an early stage at the time of sacrifice and thus only detectable with highly sensitive techniques. The cortical proteome was significantly altered by TMAO treatment in 5XFAD mice, including differential expression of proteins connected to neuronal health and amyloid processing. While many aspects of plaque pathology and neuroinflammation were unaltered, TMAO treatment did result in a significant reduction in plaque intensity; reduced plaque intensity has been associated with toxicity to surrounding neurons, which may connect with the significant reductions in neurite density and dispersion observed. Overall, our work suggests that a moderate increase in plasma TMAO worsens AD by reducing neurite density.

## ACKNOWLEDGEMENTS

This study was supported in part by grants 1R01AG070973 and T32 GM135119 of the National Institutes of Health, by the USDA National Institute of Food and Agriculture, Hatch project WIS03073, and by the University of Wisconsin-Madison School of Medicine and Public Health from the Wisconsin Partnership Program.

We thank Eugenio Vivas, Annelise Resende, and Daniela Uribe-Cano for their work in maintaining the gnotobiotic mouse breeding colony and experimental cohorts for the gnotobiotic study. We acknowledge Robert Kerby, Evan Hutchison, Sofía Murga Garrido, Connor Wilhelm, Kendra Hanslik, Kaitlyn Marino, Yajing Peng, Michael Rigby, Gonzalo Fernández-Fuente, and Luigi Puglielli for technical assistance. We appreciate the work of and services offered by the following core facilities at University of Wisconsin (UW) – Madison: Biomedical Research Model Services breeding core, the Histology Laboratory at the UW-Madison School of Veterinary Medicine, Translational Science BioCore - BioBank of the University of Wisconsin Carbone Cancer Center, Waisman Center Cellular Imaging and Analysis Core, and Optical Imaging Core.

