## Supplemental figure legends for "Trimethylamine *N*-oxide reduces neurite density and plaque intensity in a murine model of Alzheimer’s disease"

**Figure S1. Elevated plasma TMAO levels do not alter the cortex proteome of WT mice.** (A) Plasma TMAO levels were significantly increased in animals supplemented with TMAO. (B) Fold change (FC) of protein abundances between TMAO-treated WT mice *vs.* the control. Volcano plot of log_2_ fold change of TMAO over untreated mice *vs.* log_10_ q-value. The horizontal dashed line represents q = 0.05 and the vertical dashed lines represent FC = |1.5|; proteins with a q-value < 0.05 and FC > |1.5| are shown as orange circles with a black border.

**Figure S2. Gnotobiotic mice colonized with TMA(O)-producing community exhibit a trend towards lowered amyloid plaque burden in cortex.** Germ-Free (GF) mice were colonized with either a choline-consuming/TMAO-producing (CC^+^) or non-choline-consuming/non-TMAO-producing (CC^-^) community between 5-7 weeks of age and placed on a 1% choline diet (Envigo TD.140179); mice were sacrificed at 32 weeks of age (A). (B) Circulating plasma TMAO levels were significantly higher in CC^+^ mice. (C) Two brain sections per animal were stained for plaques (6E10, brown) and counterstained with hematoxylin (purple). (D) The percent area positive for 6E10 in the cortex and the hippocampus was quantified using FIJI. Data are shown as the mean ± SEM (n=4 CC^-^, n=4 CC^+^, two sample t-test). ** denotes p < 0.01.
