## Supplementary figures and images for "Trimethylamine *N*-oxide reduces neurite density and plaque intensity in a murine model of Alzheimer’s disease"

### Supplemental Figure 1

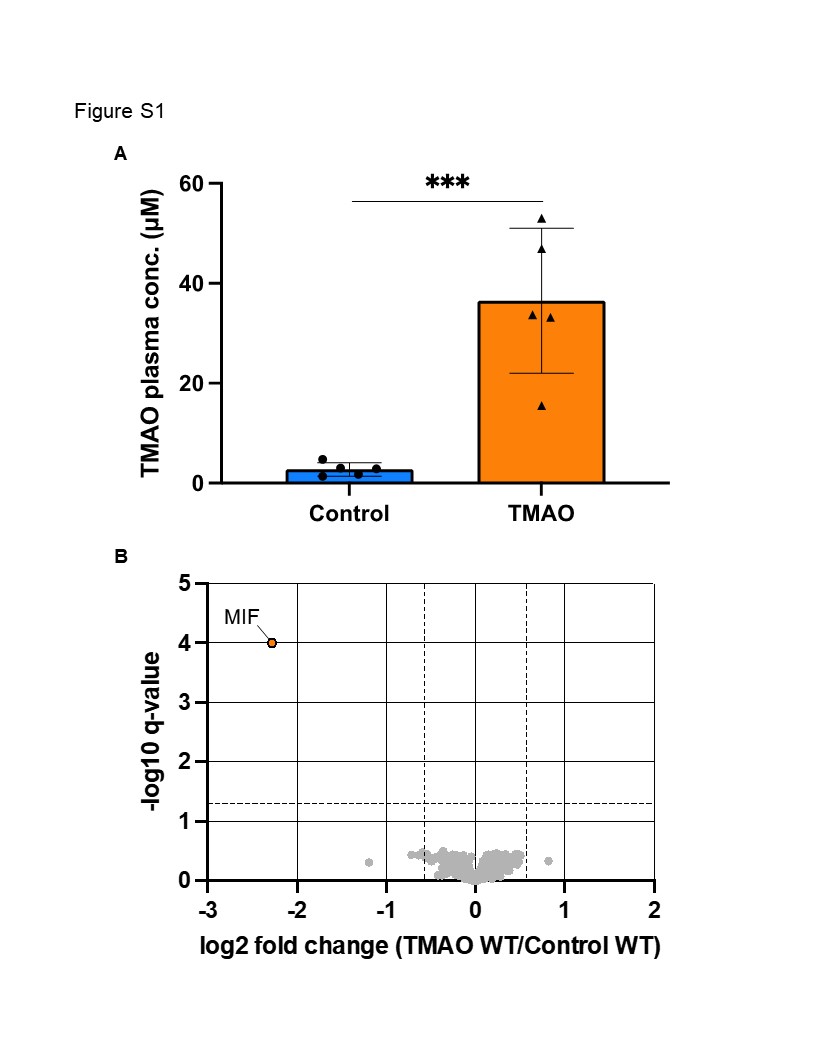

### Supplemental Figure 2

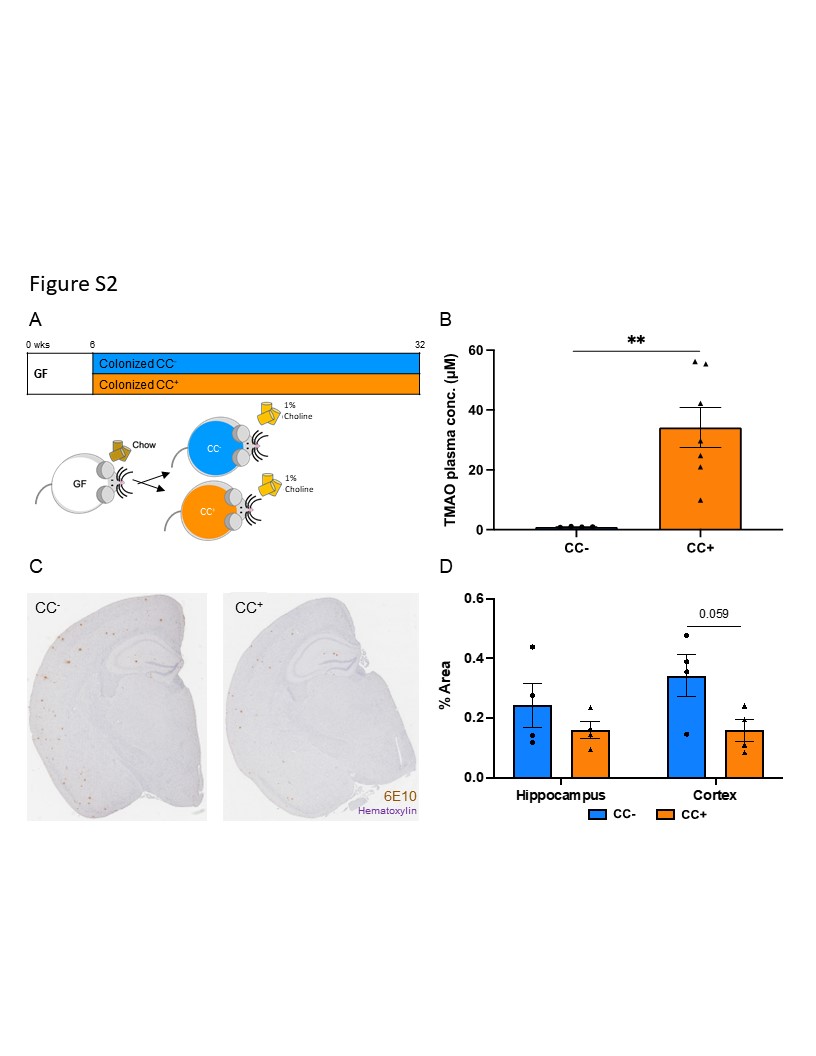
